# Development of a High-Throughput Minimum Inhibitory Concentration (HT-MIC) Testing Workflow

**DOI:** 10.1101/2022.10.21.513098

**Authors:** Suman Tiwari, Oliver Nizet, Nicholas Dillon

## Abstract

The roots of the minimum inhibitory concentration (MIC) determination go back to the early 1900s. Since then, the test has undergone modifications and advancements in an effort to increase its dependability and accuracy. Although biological investigations use an ever-increasing number of samples, complicated processes and human error sometimes result in poor data quality, which makes it challenging to replicate scientific conclusions. The automation of the few manual steps using protocols decipherable by machine can ease some of the procedural difficulties. Originally relying on manual pipetting and human vision to determine the results, modern broth dilution MIC testing procedures have incorporated microplate readers to enhance sample analysis. However, current MIC testing procedures are unable to simultaneously evaluate a large number of samples efficiently. Here, we have created a workflow using the Opentrons OT-2 robot to enable high-throughput MIC testing. We have further optimized the analysis by incorporating Python programming for MIC assignment to streamline the automation. In this workflow, we performed MIC tests on four different strains, three replicates per strain, and analyzed a total of 1,152 wells. Comparing our our workflow to a conventional plate MIC procedure, we find that the HT-MIC method is 630% faster while simulataneously boasting a 100% accuracy. Our high-throughput MIC workflow can be applied in both academic and clinical settings since it is faster, more efficient, and more accurate than many conventional methods.

## Introduction

Alexander Fleming first reported the inhibitory effect of antibiotics against staphylococci in 1929 by serially diluting penicillin in nutrient broth. He observed that the opacity of the broth, or the culture’s pH, could be used to detect the inhibition of a bacterial suspension (1, 2). Through expanding this technique he made it possible to gauge the inhibitory potency of antibiotics against various bacterial species. Fleming’s early work has been described as a forerunner of contemporary minimum inhibitory concentration (MIC) methodology. A MIC is defined as the lowest concentration of an antibacterial agent expressed in mg/mL (g/L) which, under strictly controlled *in vitro* conditions, prevents visible growth of the test strain of an organism (3).

Antibiotic susceptibility testing using standardized disk diffusion was introduced by Bauer and Kirby’s experiments in 1956. In this method, the isolated bacterial colony is selected, suspended into growth media, and standardized through a turbidity test. The standardized bacterial suspension is inoculated onto a solidified agar plate and an antibiotic-treated paper disk is added. The antibiotic on the disc is allowed to permeate through the agar overnight, forming a zone of killed bacteria (4). Whilst antimicrobial activity methods based on agar diffusion have clear benefits in terms of efficiency and cost, they are not without drawbacks, and the validity and appropriateness of agar diffusion methods for accurately quantifying antimicrobial activity has been questioned (5, 6). With the advancement of technology, MIC values were later determined by broth dilution using a microplate reader to reduce the dependance on human observation.

Microplate-readers are instruments designed to measure the absorbance, fluorescence, or luminescence of samples in microtitre plates and are used for a variety of applications (7). The microtiter plates, first introduced in the 1970s, normally have 96 wells, each of which has a volume of 100–200 μL for MIC experiments (8). Their primary benefits include low sample volumes, high experimental throughput, and simplicity of usage.

In the 1980s the Clinical and Laboratory Standards Institute (CLSI) consolidated the methods and standards for MIC determination for clinical usage (9). Large sets of MIC sampling, including multiple strains and antibiotics, are time-consuming using the standardized approaches as they are currently not automated. We require new high-throughput sample handling methods as big data usage is becoming more commonplace. Recent developments in information technology and robots have optimized manual labor operations and could now be used to modernize and streamline MIC testing (10).

We have developed a high-throughput MIC (HT-MIC) testing workflow to overcome the shortcomings of previous approaches. HT-MIC uses a robot to create the testing plates, determines the inhibition using a microplate reader, and uses Python programming to assign MIC values that are largely free of human error. The HT-MIC workflow uses an Opentrons OT-2 robot (Opentrons Inc., New York, NY, USA), a benchtop liquid handling tool that can transfer liquids using disposable tips and electronic pipettes (11). The OT-2 is one of many automated liquid handling systems now on the market, but it stands out due to its inexpensive cost, which should make it affordable for many laboratories (12). A wide variety of industrial processes have seen improvements in their production rates, efficiency, and quality thanks to the gradual incorporation of automation into workflows (13). We propose the HT-MIC method is faster, more efficient, and more accurate than conventional methods.

## Materials and equipment

### Opentrons Robot

The workflow mentioned in this paper utilizes the Opentrons OT-2 robot with removable P300 multichannel Gen1 pipette mounted on the right side and P50 single channel Gen1 pipette (not used in the protocol) mounted on the left side of the robot.

### Medium

The most often used medium for broth dilution techniques globally is Cation Adjusted Mueller-Hinton broth (CA-MHB) which is used in this workflow as well. In general, it has few antagonists and promotes healthy growth of the majority of nonfastidious pathogens (14).

### Bacterial strains and antibiotics

*Acinetobacter baumannii* strains AB 19606, AB 1605, AB 1710, and AB 1789 were obtained from the American Type Culture Collection (ATCC) and stored at - 80°C in 20% glycerol (Biobasic) and 80% CA-MHB (Difco) (15).

Concentrated stocks of azithromycin (Fresenius Kabi) and ciprofloxacin (Sigma) were prepared at 100□mg/mL in DI H_2_0. Doxycycline (Fresenius Kabi), vancomycin (Mylan), and meropenem (Hospira) were prepared at 50 mg/mL in DI H_2_0. Minocycline (Melinta) at 20□mg/mL, tigecycline (Fresenius Kabi) at 0.45 mg/mL, and levofloxacin (Sigma) at 1.44 mg/mL were all also prepared in DI H_2_0. Fresh 100× antibiotic experimental stocks were made at the desired concentration prior to the start of each experiment (Fig. 1A).

**Fig 1).**
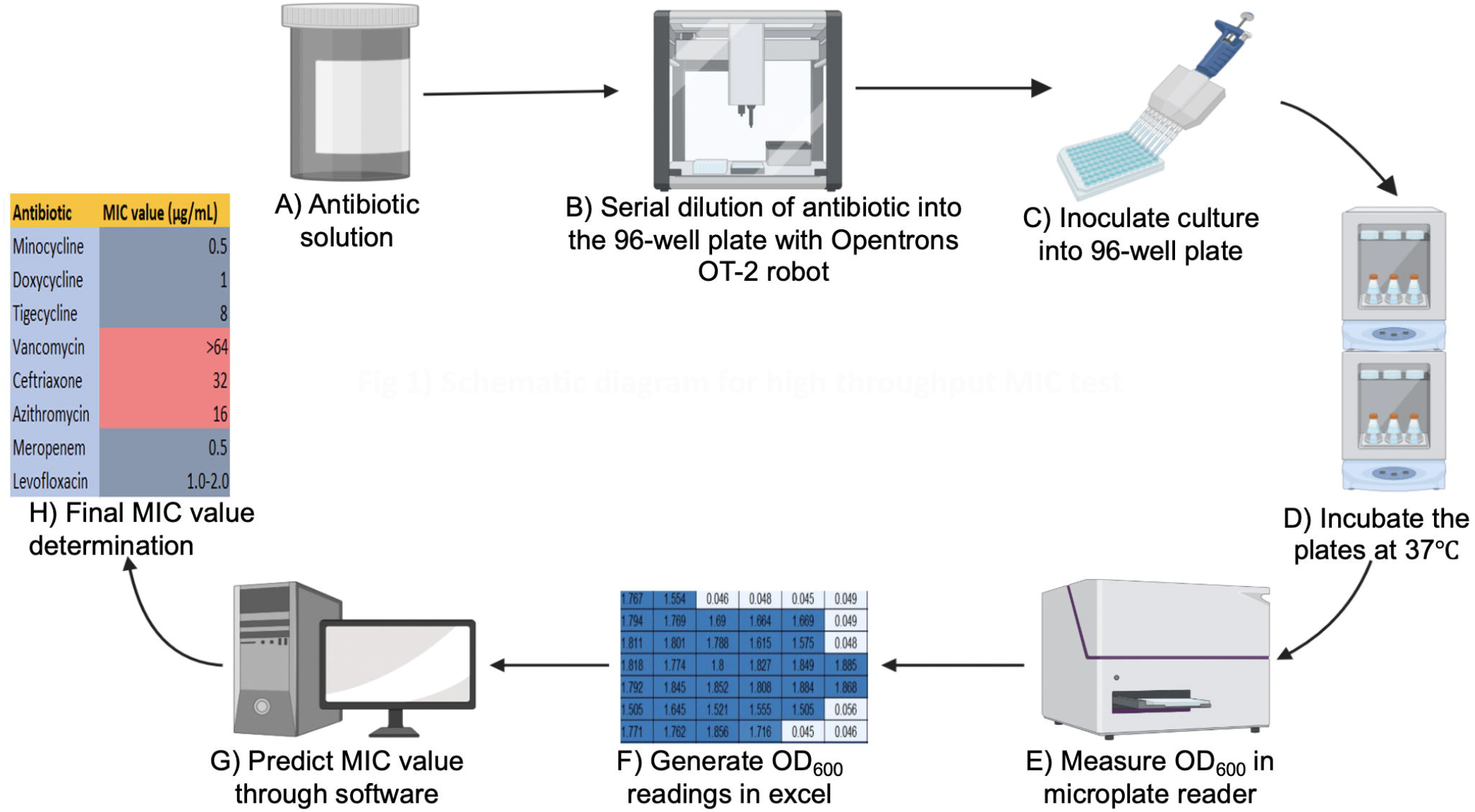
Schematic diagram for high throughput MIC test

## Methods

### Opentrons workflow

Figure 1 illustrates the workflow for our HT-MIC testing method using multiple modules on the Opentrons OT-2 robot. The protocol for the use of the OT-2 robot was created using the Opentrons protocol designer beta program. Before using, the Opentrons robotic platform was calibrated. The idea is to manually put the liquid-handling arm in the predetermined position at each slot (such as 96-well plates, a tip box, the garbage, etc.) to enable the robot to remember that position. (15). The workflow begins with the pre-processing module to generate an antibiotic master plate containing serially diluted working stocks of each antibiotic. In the first step of this module CA-MHB is added to each well of columns 1-11 (150 μL) and to column 12 (270 μL) to facilitate serial dilutions. Each 100x antibiotic stock is then added to column 12 of the 96-well plate through diluting 30 μL of the 100x stock into 270 μL of media to create a final 10x working stock. Well contents are mixed, and the pre-processing module pipettes 150 μL from column 12 to column 11 to start the 2-fold dilution scheme. 2-fold serial dilutions continue from column 12 until column 2 with the 300 μL pipette tips (Opentrons) discarded and reloaded after dispensing into column 9, column 6 and Column 2. Using this layout scheme each row contains 11 concentrations of each antibiotic arrayed in 2-fold increments. Antibiotics are intentionally omitted from column 1 to create a no drug control (Fig. 2). In the next step, the antibiotic master plate is used to create 5 identical MIC test plates. The Opentrons deck state for five 96-well MIC test plates is shown utilizing 300 μL pipette tips (Opentrons) in deck 1 and 2, the master plate in deck 3, the test plates in decks 4-8 (Fig. 3) and 90 mL well reservoir (Axygen) in deck 9. 30 μl of antibiotic solution from each well of the master plate is replica plated into each of the five 96-well MIC test plates. These steps can be repeated multiple times using the protocol until the desired quantity of 96-well plates is made. The code for the pre-process module for the Opentrons OT-2 robot is provided in the GitHub repository under “antibiotics.json” file name. JavaScript Object Notation (JSON) documents are both machine- and human-readable, and storing the metadata in the JSON format preserves the adaptability of the created metadata with various research data management tools and processes (16).

**Fig 2).**
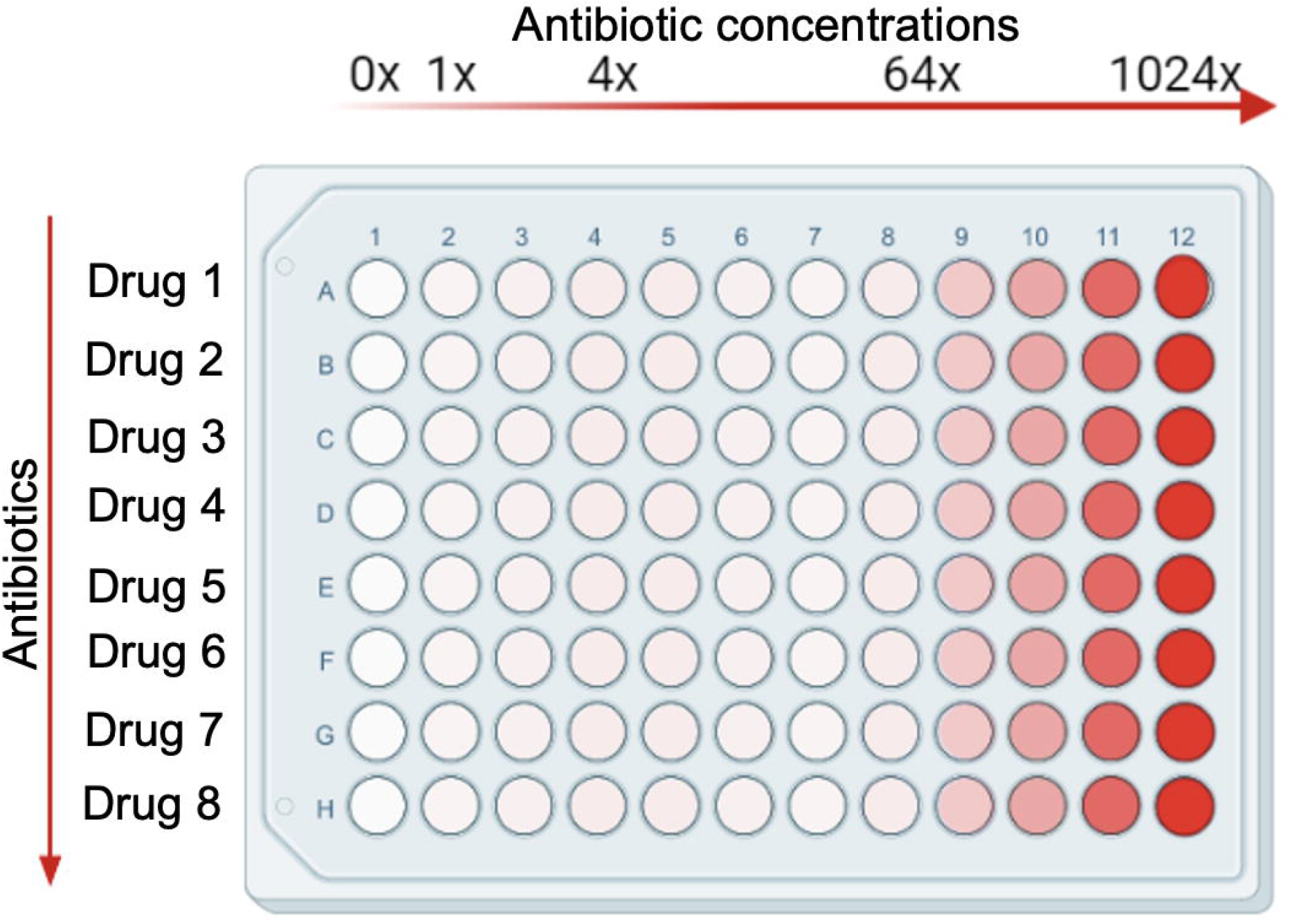
2-fold concentration increment of antibiotics in 96-well plate

**Fig 3).**
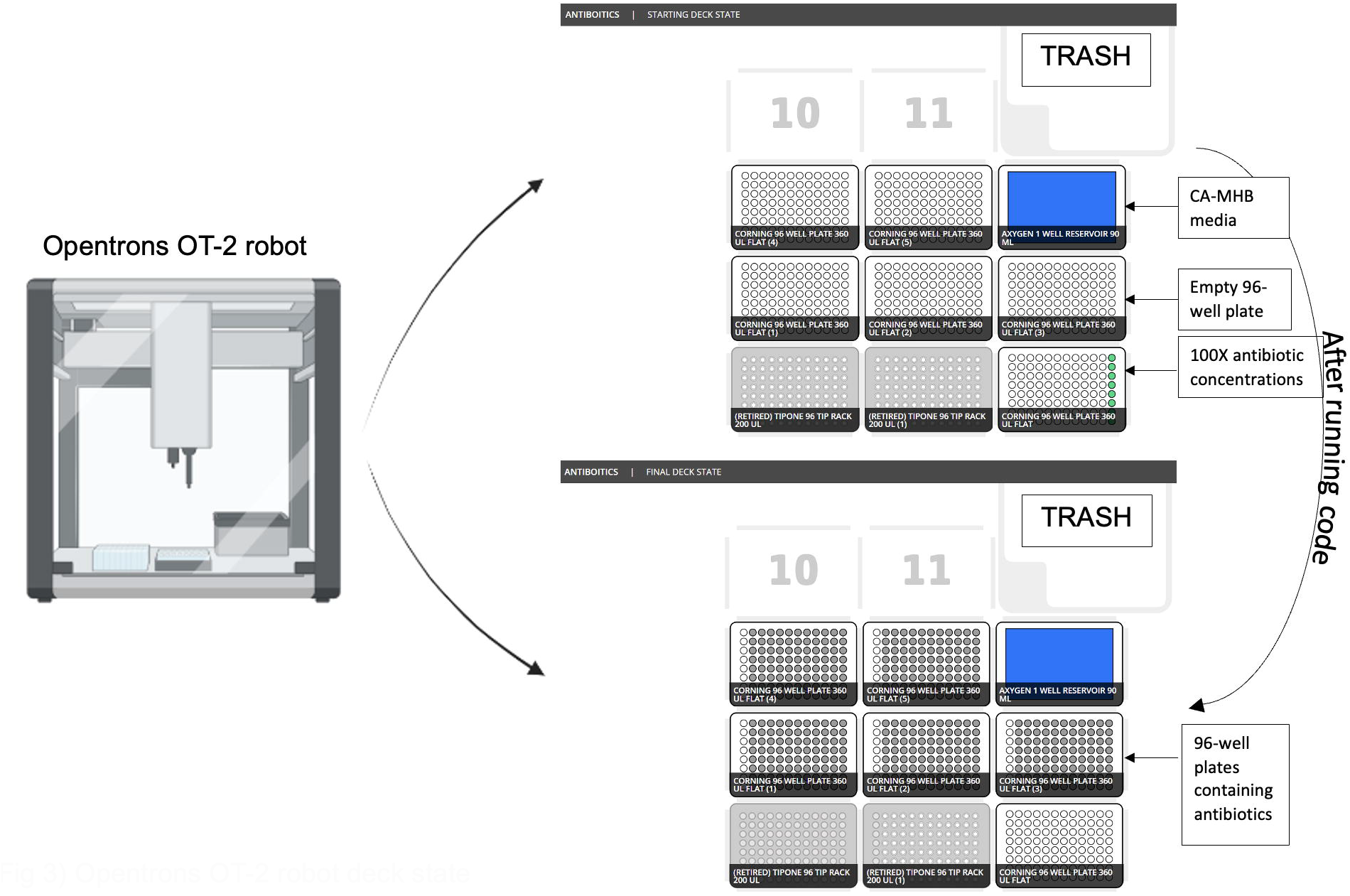
Opentrons OT-2 robot deck state

### MIC testing

Each independent replicate culture was grown overnight at 37°C shaking on an orbital shaker (Innova) at 200 rpm in CA-MHB (+ 10 mg/L MgCl_2_ (Fisker) and 20 mg/L CaCl_2_ (Sigma) to stationary phase. The optical density (OD) of a bacterial cell culture can be determined spectrophotometrically, which is a practical and widely used approach to assess its growth stage (17). Fifty μL of overnight cultures was inoculated into fresh CA-MHB the day of the experiment and grown to mid-log phase (OD_600_=∽0.4, ∽1*10^8^ CFU/mL) at 37°C and 200 rpm (15). Mid-log phase cultures were diluted to ∽ 1*10^5^ CFU/mL (OD_600_=∽0.002) and 270 μL of cultures were added to 96-well (Costar) plates that contained 30 μL antibiotics dispensed using our HT-MIC Opentrons workflow protocol (Fig. 1C) provided in the Github repository under “MIC test.json” file name. In this protocol, there is 300 μL pipette tips (Opentrons) in deck 1 and 2, 90mL well reservoir (Axygen) containing cultures in deck 3, and stock antibiotic plates in deck 4 and 5. Plates containing both bacterial cultures and antibiotics were placed into the orbital shaker and grown for approximately 20 hrs shaking at 200 rpm at 37°C (Fig 1D). After incubation, the 96-well plates were removed and the OD_600_ of each well was measured on a 96-well plate reader (Synergy H1) (Fig 1E). Measured OD_600_ values were exported into excel for MIC determination (Fig 1F). This protocol is used to determine high confidence MIC_90_ values. The MIC_90_ is defined as the lowest concentration of an antibiotic that inhibits ≥10 % of the bacterial growth found in the no-antibiotic control (18).

### Automated MIC determination

The automated MIC determination is accomplished using a Python program provided in GitHub repository under “MIC-PythonCode.docx” file name. The program input is an Excel file comprised of two sheets provided in GitHub repository under “example_input data.xlsx” file name. The first sheet includes the measured OD_600_ values for three different 96-well plates, each sharing the same antibiotics and concentration ranges. Each row in the spreadsheet corresponds to one of the eight antibiotics, and the 36 columns correspond to the OD_600_ readings repeated in triplicate for each of 12 concentrations. The second sheet possesses metadata detailing the concentrations of antibiotic corresponding to each of the OD_600_ values on the first sheet. The program first reads the Excel file and formats the incoming data, then for each antibiotic searches the metadata (2nd sheet) to locate which columns have the same concentrations, then groups the indices. The program subsequently uses those indices to group the OD_600_ readings (1st sheet) by concentration and antibiotic.

For each antibiotic, the program goes starts at the lowest concentration and checks if all three OD_600_ readings are above 0.15.

- If all three readings are above 0.15, it moves to check the next lowest concentration.
- If two of the readings are above 0.15, it determines if the third is at least above 0.135. If the third is above 0.135, it moves to check the next lowest concentration. If the third is below 0.135, the current concentration is the MIC
- If one of the readings is above 0.15, it checks if that reading is below
- 0.165. If that reading is below 0.165, the current concentration is the MIC. If that reading is above 0.165, it moves to check the next lowest concentration.
- If none of the readings are above 0.15, then the current concentration is the MIC

After checking all the concentrations in this manner, an MIC is either determined or the MIC is labeled to be greater than the highest concentration. The program finally outputs an 8×2 spreadsheet as a .csv file with one column listing every antibiotic tested and the other column showing the corresponding MIC. Program run time is nearly instantaneous.

## Results

To assess the accuracy of the high-throughput MIC prediction method we did a side-by-side comparison of the individual assigned MIC_90_ values vs. those determined by the newly developed prediction model from the HT-MIC workflow. The predicted MIC value from the HT-MIC workflow accurately identified the MIC_90_ values determined from the individual assigned method without any discrepancies (Table 1). This absolute concordance shows that the prediction model is accurate and in line with the standard method of assigning MIC values. All outputs of 1,152 wells given by microplate reader is provided in the Github repository under “MIC-4 strains list.xlsx” file name.

**Table 1:** Comparison of MIC values of microplate reader vs software

| Strains<br>Antibiotics | AB 1605 | AB 1710 | AB 1789 | AB 19606 |
| --- | --- | --- | --- | --- |
| Minocycline | 2.0 | 1.0 | 2.0 | 0.5 |
| Doxycycline | 32.0 | 16.0 | 16.0 | 1.0 |
| Tigecycline | 16.0 | 16.0 | 16.0 | 8.0 |
| Vancomycin | 64.0 | >64.0 | >64.0 | >64.0 |
| Ceftriaxone | >64.0 | >64.0 | >64.0 | 32.0 |
| Azithromycin | 16.0 | 16.0-32.0 | 16.0 | 16.0 |
| Meropenem | 8.0 | 8.0 | 8.0 | 0.5 |
| Levofloxacin | >64.0 | >64.0 | >64.0 | 1.0-2.0 |

MIC value in agreement with both techniques MIC not in agreement

Having demonstrated the accuracy of the method, we next compared the efficiency of the HT-MIC method to that of the standard MIC 96-well plate testing scheme. The HT-MIC workflow is much more efficient, and faster than the standard MIC method (Fig. 4). To quantify this we compared the amount of time needed to determine the MIC_90_ for 8 antibiotics (each examined at 11 different concentrations) in 4 bacterial strains in triplicate yielding 96 determined MIC_90_ values. Using standard MIC protocols, creating the 30 antibiotic plates needed for the MIC testing takes ∽11 hrs. Using the HT-MIC workflow, it took only ∽2 h to make the same set of 30 plates, a savings of ∽9 h (Fig. 4). Similarly, the inoculation of bacterial took ∽3 h for 30 plates when done through standard methods while the automated process accomplished it in less than 30 min (Fig. 4). Moreover, the MIC prediction from automated code is instantaneous compared to the ∽1 hr analyzing the readings manually. Overall, MIC determination takes ∽11 hrs to complete for 30 plates using the standard MIC method, while its takes only ∽1.5 hrs for the same 30 plates if done by HT-MIC method. We determined that the HT-MIC method is ∽630% faster for determining MIC values compared to standard MIC testing. Not only is the HT-MIC method faster, but it is also more efficient as it negates human pipetting error as the robot has significantly less errors.

**Fig 4).**
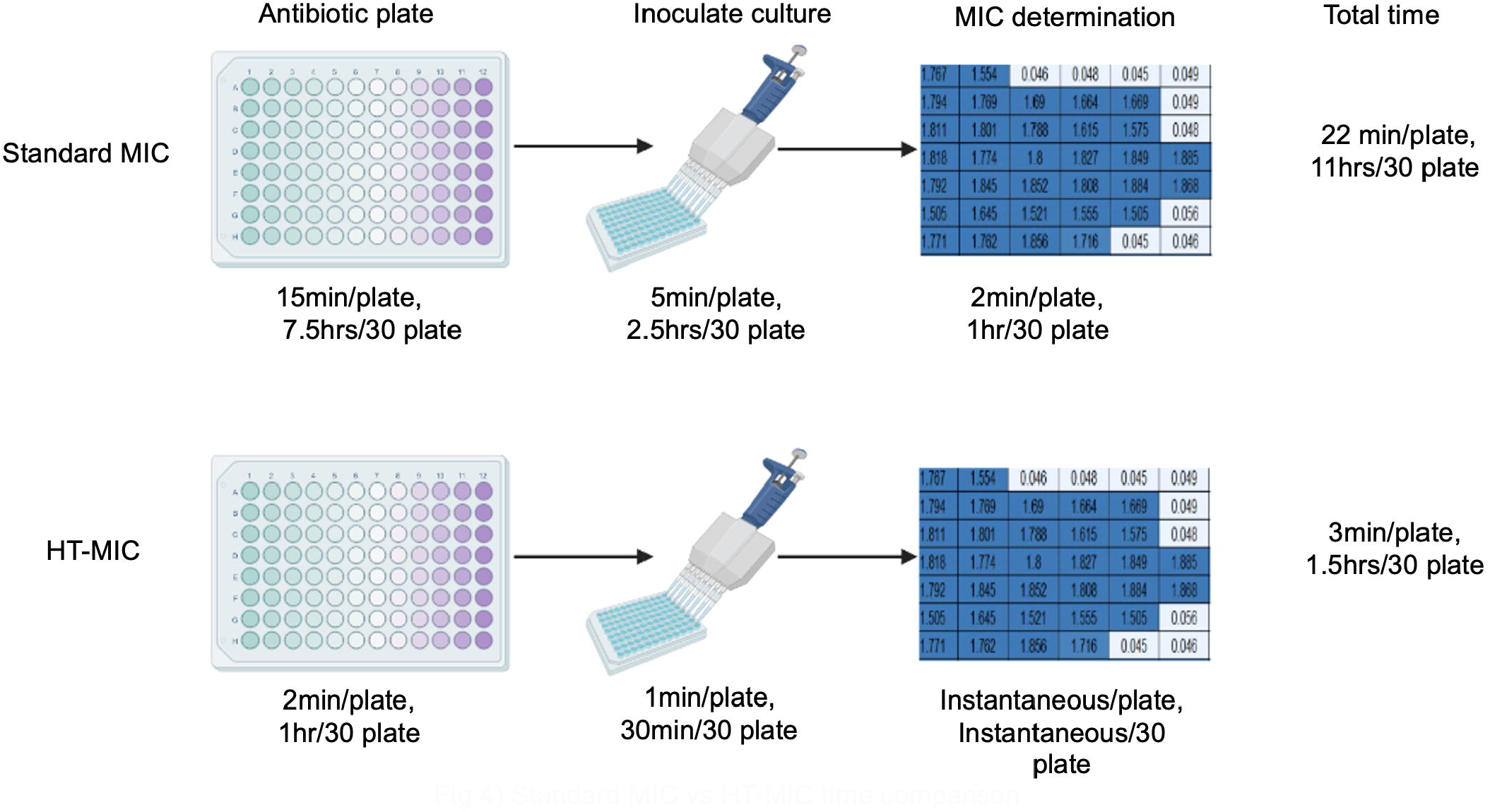
Standard MIC vs HT-MIC time comparison

## Discussion

The HT-MIC workflow was demonstrated to be a more efficient and accurate method for measuring MIC_90_ values when compared to standard methods. This method utilizes an Opentrons OT-2 robot with a customized workflow to permit high-throughput antibiotic sensitivity experiments yielding more data with a higher degree of accuracy. The Opentrons robot’s comparatively simple setup and operation (programmable via a Python API or via a graphical user interface) allows for flexible use by non-specialized workers. Hopefully, the straightforward sharing of methods will make it simple for other labs to check and validate the work of others (19).

The protocols used in the HT-MIC workflow are accessible through the GitHub repository provided in this paper. The protocol can be downloaded and amended for use with the Opentrons OT-2 robot to fit any specific experimental requirements. The protocol can be modified in the Opentrons protocol designer beta program allowing for the editing of steps like changing the volumes, dilution, or transferring of solution if desired.

This automated time efficient method is applicable to not only researchers in academic setting but could also be incorporated into clinical settings. Clinical microbiology labs often deal with a large influx of patient samples necessitating efficient testing processes. Using this method, sensitivity testing can be conducted with different antibiotics for multiple samples simultaneously. The automated method is hassle free and can be carried out with high accuracy.

The protocol provided in the GitHub repository could be improved even further to decrease the experimental time and increase the data generated. Only 5 antibiotic stock plates are made using the provided protocol. This limitation is due to the use of standard 96-well plates for the entirety of the workflow. However, efficiency could be improved through using deep well plates for the antibiotic stock plates instead of 96-well plates as larger volumes can be made. The same master antibiotic plate could then be used to make 40 antibiotic testing plates instead of 5. Using deep well plates would save even more time and will make the HT-MIC workflow even more efficient.

The link to the GitHub repository is provided here: https://github.com/suman921/HT-MIC

## References

1. Fleming A. On the antibacterial action of cultures of a penicillium, with special reference to heir use in the isolation of ‘B. influenzae’ 1929.

2. Wheat PF. History and development of antimicrobial susceptibility testing methodology. J Antimicrob Chemother. 2001;48 Suppl 1:1–4.

3. Kowalska-Krochmal B, Dudek-Wicher R. The Minimum Inhibitory Concentration of Antibiotics: Methods, Interpretation, Clinical Relevance. Pathogens. 2021;10(2).

4. Khan ZA, Siddiqui MF, Park S. Current and Emerging Methods of Antibiotic Susceptibility Testing. Diagnostics (Basel). 2019;9(2).

5. Bonev B, Hooper J, Parisot J. Principles of assessing bacterial susceptibility to antibiotics using the agar diffusion method. J Antimicrob Chemother. 2008;61(6):1295–301.

6. Eloff JN. Avoiding pitfalls in determining antimicrobial activity of plant extracts and publishing the results. BMC Complement Altern Med. 2019;19(1):106.

7. Thompson JE. Low-Cost Microplate Reader with 3D Printed Parts for under 500 USD. Sensors (Basel). 2022;22(9).

8. Ashour MB, Gee SJ, Hammock BD. Use of a 96-well microplate reader for measuring routine enzyme activities. Anal Biochem. 1987;166(2):353–60.

9. Wikler MA. Methods for dilution antimicrobial susceptibility tests for bacteria that grow aerobically: approved standard. Clsi (Nccls). 2006;26:M7–A.

10. Holland I, Davies JA. Automation in the Life Science Research Laboratory. Front Bioeng Biotechnol. 2020;8:571777.

11. Tenhaef N, Stella R, Frunzke J, Noack S. Automated Rational Strain Construction Based on High-Throughput Conjugation. ACS Synth Biol. 2021;10(3):589–99.

12. Liang Y, Acor H, McCown MA, Nwosu AJ, Boekweg H, Axtell NB, et al. Fully Automated Sample Processing and Analysis Workflow for Low-Input Proteome Profiling. Anal Chem. 2021;93(3):1658–66.

13. Hitomi K. Automation — its concept and a short history. Technovation. 1994;14(2):121–8.

14. Microbiology ECfASTotESoC, Diseases I. Determination of minimum inhibitory concentrations (MICs) of antibacterial agents by broth dilution. Clinical Microbiology and Infection. 2003;9(8):ix–xv.

15. Dillon N, Holland M, Tsunemoto H, Hancock B, Cornax I, Pogliano J, et al. Surprising synergy of dual translation inhibition vs. Acinetobacter baumannii and other multidrug-resistant bacterial pathogens. EBioMedicine. 2019;46:193–201.

16. Chaerony Siffa I, Schäfer J, Becker MM. Adamant: a JSON schema-based metadata editor for research data management workflows. F1000Res. 2022;11:475.

17. Meyers A, Furtmann C, Jose J. Direct optical density determination of bacterial cultures in microplates for high-throughput screening applications. Enzyme Microb Technol. 2018;118:1–5.

18. Liang S, Kinghorn AB, Voliotis M, Prague JK, Veldhuis JD, Tsaneva-Atanasova K, et al. Measuring luteinising hormone pulsatility with a robotic aptamer-enabled electrochemical reader. Nat Commun. 2019;10(1):852.

19. Jessop-Fabre MM, Sonnenschein N. Improving Reproducibility in Synthetic Biology. Front Bioeng Biotechnol. 2019;7:18.

